# Genotyping-by-sequencing based cytoplasmic markers underpin population structural variation of perennial ryegrass

**DOI:** 10.1101/610543

**Authors:** Mingshu Cao, Marty Faville, Jeanne Jacobs, Marcelo Carena

## Abstract

Chloroplast and mitochondrial genomes provide unique information in studying plant populations because cytoplasmic genes exhibit a different mode of inheritance and a different rate of gene mutation compared to nuclear genes. Despite this, cytoplasmic genomic contributions to plant population performance are largely unexplored because few methods are available to characterize and evaluate cytoplasmic genome-wide variations. Here we have developed cytoplasmic markers based on genotyping-by-sequencing (GBS), which enable us to characterize thousands of samples, to survey gene variants across cytoplasmic genomes, and to monitor within-population variations of chloroplast or mitochondrial origin. Using these cytoplasmic genome-wide markers we have found that within-population differentiations are evident in ryegrass (*Lolium perenne*), beyond the explanation of nuclear markers. Moreover, chloroplast and mitochondrial variations exhibit different patterns, with mitochondrial markers more readily reflecting the maternal origins. Application of GBS-based cytoplasmic markers should facilitate quantifying the contribution of cytoplasmic inheritance to plant performance through selective breeding or under natural selection pressure.

## Introduction

Plant cytoplasmic genomes of chloroplasts and mitochondria provide unique information for studying population diversity because genes of cytoplasmic origin exhibit a different mode of inheritance and a different rate of gene mutation compared to nuclear genes (Wolfe et al. 1987). Thanks to their relatively small size complete mitochondrial or chloroplast genomes have been sequenced for 3,015 plant species, according to the NCBI organelle genome databases (https://www.ncbi.nlm.nih.gov/genome/organelle/ accessed at April 2019). Organelle genome information has contributed to our understanding of phylogenetic relationships or evolution among many plant species. That information was based on variations estimated from complete chloroplast DNA (cpDNA), mitochondrial DNA (mtDNA), or targeted genes residing in cytoplasmic genomes (Hiesel et al. 1994; Dobler et al. 2014; Daniell et al. 2016). At the level of microevolution (i.e. from the perspective of plant improvement), cpDNA and mtDNA sequence-informed genetic markers, such as simple sequence repeat (SSR) markers, have been used to study plant domestication, to validate the parental origin of commercially important varieties, and to characterize genetic variation among germplasms or cultivars (Galtier et al. 2009; Daniell et al. 2016). Nonetheless, all such studies were based on markers derived from a few genes or repeat regions, and on a limited number of individual plants due to the limited capability of genotyping methods.

In breeding for outcrossing plant species, selections are often carried out at the population or family level. Accurately monitoring gene allele frequency allows desirable genotypes to predominate in selected populations, and this calls for efficient genotyping tools that are deployable at a large scale. Genotyping-by-sequencing (GBS) (Elshire et al. 2011), due to its multiplexing and use of restriction enzyme(s) to reduce genome complexity, represents one of the productive and affordable options for characterizing a large number of samples to meet the demand for genetic improvement of populations. We have applied GBS to characterize perennial ryegrass (*Lolium perenne* L.) populations based on nuclear DNA (nuDNA) derived single nucleotide polymorphism (SNP) markers (Faville et al. 2018). Given a massive amount of sequencing tags sampled from each individual plant we anticipate that a certain number of tags will be derived from cpDNA and mtDNA, thus could be utilized for cytoplasmic SNP discovery. Organellar DNA is expected to be contained in routine whole genomic DNA isolations (Islam et al. 2013), and organelle genomes could even be assembled from whole genome data (Dierckxsens et al. 2017).

In perennial ryegrass the mitochondrial genome (Islam et al. 2013) and chloroplast genome (Diekmann et al. 2009; Hand et al. 2013) have been assembled. In those reports perennial ryegrass chloroplast and mitochondrial genomes were studied in comparison with that from more or less related species, the purposes of which were to gain insights into phylogenetic relationships. More relevant to the context of our investigation here is whether these types of variations can be harnessed at the level of cultivar or germplasm (of the same species). Polymorphism was indeed found among a few individual *L. perenne* plants from the same population based on cpDNA-derived microsatellite markers (Diekmann et al. 2012), or based on SNPs detected from chloroplast sequencing reads (Diekmann et al. 2009). However, no systematic information is available on the extent of cpDNA and mtDNA variation occur within ryegrass cultivars or among cultivars. With its scalability and free of ascertainment bias GBS may provide an opportunity to assess such genetic variation within and among populations.

With an aim to explore and exploit that opportunity we have developed a computational workflow for haploid SNP discovery from mapping existing GBS reads to cytoplasmic reference genomes (cpDNA and mtDNA); we quantified the genetic similarity of individual plants and assess genetic polymorphism within ryegrass breeding populations using distance metrics. Potential uses of GBS-based cytoplasmic markers for breeding outcrossing and perennial plant species are discussed.

## Results

### Identification and characterization of cytoplasmic markers from ApeKI GBS libraries

We used GBS data derived from two ryegrass breeding populations ‘P96’ (112 plants) and ‘P127’ (120 plants). Four controls samples from the same DNA extraction of a single individual were used to assess the reproducibility of SNP discovery.

A total of 5,115,152 quality tags were generated from the 236 samples. Only 8,807 unique tags were mapped onto the cytoplasmic reference genomes, among which 6,056 and 2,751 tags were mapped to cpDNA and mtDNA, respectively. 1,071 raw haploid SNPs were obtained using samtools and bcftools. Fewer SNPs (197) were detected with cpDNA as the reference than the number of SNPs (874) detected with mtDNA. The two sets of SNPs are hereafter referred to as cpSNPs and mtSNPs.

The control samples, although of the same DNA extraction, were subjected to four independent GBS library preparations and sequenced in different batches (sequencing flowcells and lanes). 95.3% (1021 out of 1071) SNPs were consistently identified (sd = 0, n=4) among the controls, demonstrating the reliability of SNP calling. The 50 inconsistent SNPs appeared to be random in the controls, and its variation was not associated with the low read depth (mean of read depth of the 50 SNPs = 644.7). We thus removed these inconsistent SNPs. Among the retained SNPs, 649 were called with the constant value of 1 across both populations, being 1 means the site genotype has a different nucleotide from the corresponding reference site (see Methods). Although different from the reference the monomorphism among the samples under the study offers no information we therefore removed them from further analysis. As a result, 372 SNPs (100 cpSNPs and 272 mtSNPs) were retained for statistical analysis.

Phi correlation coefficient (ϕ) was applied to measure genetic relationships of the samples based on the retained SNPs. Multidimensional scaling (MDS) analysis performed on the distance metric (1-ϕ) showed somewhat complex relationships between populations and within each population, based on both cpSNPs and mtSNPs (Fig. 1).

**Fig. 1.**
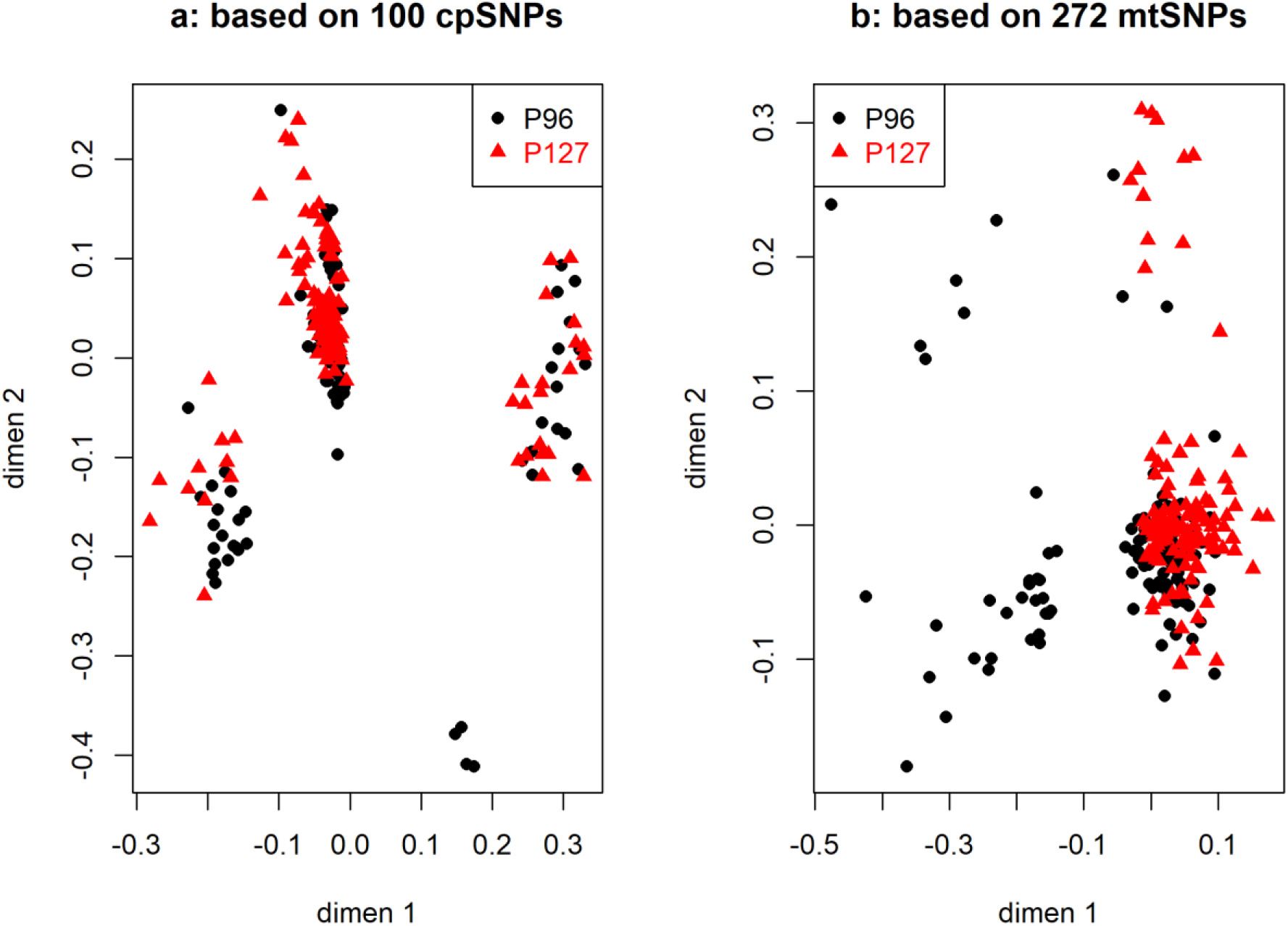
Multidimensional scaling clustering of samples from perennial ryegrass populations P96 (n=112) and P127 (n=120), based on cpSNPs (a) and mtSNPs (b).

Substructures within both ‘P96’ and ‘P127’ populations were clearly revealed based on cpSNPs (Fig. 1a). It was unexpected to see that the sample clustering did not reflect the two populations, with three clusters nonetheless observed only within the respective population (Fig. 1a). Sanity checking via principal component analysis confirmed the substructures were not associated with the GBS libraries, sequencing batches, nor the amount of tags per sample. On the other hand, based on mtSNPs ‘P96’ largely formed two clusters across the dimension 1, with discernible within-population differentiations (Fig. 1b).

Excluding possible confounding technical factors (such as the library and sequencing batches) the substructure revealed should be attributed to the intrinsic SNP variations. A close inspection of the variation patterns of cpSNPs among the samples confirmed that this was the case. There were 3 clusters of plants within ‘P96’ (Fig. 1a and Supplementary Fig.1), where, as shown in Fig. 2 (limited view due to the space), plants 35, 38, 44, 48, 58, 60, 63 belonged to the same class as the same pattern of SNP variation was presented.

**Fig. 2.**
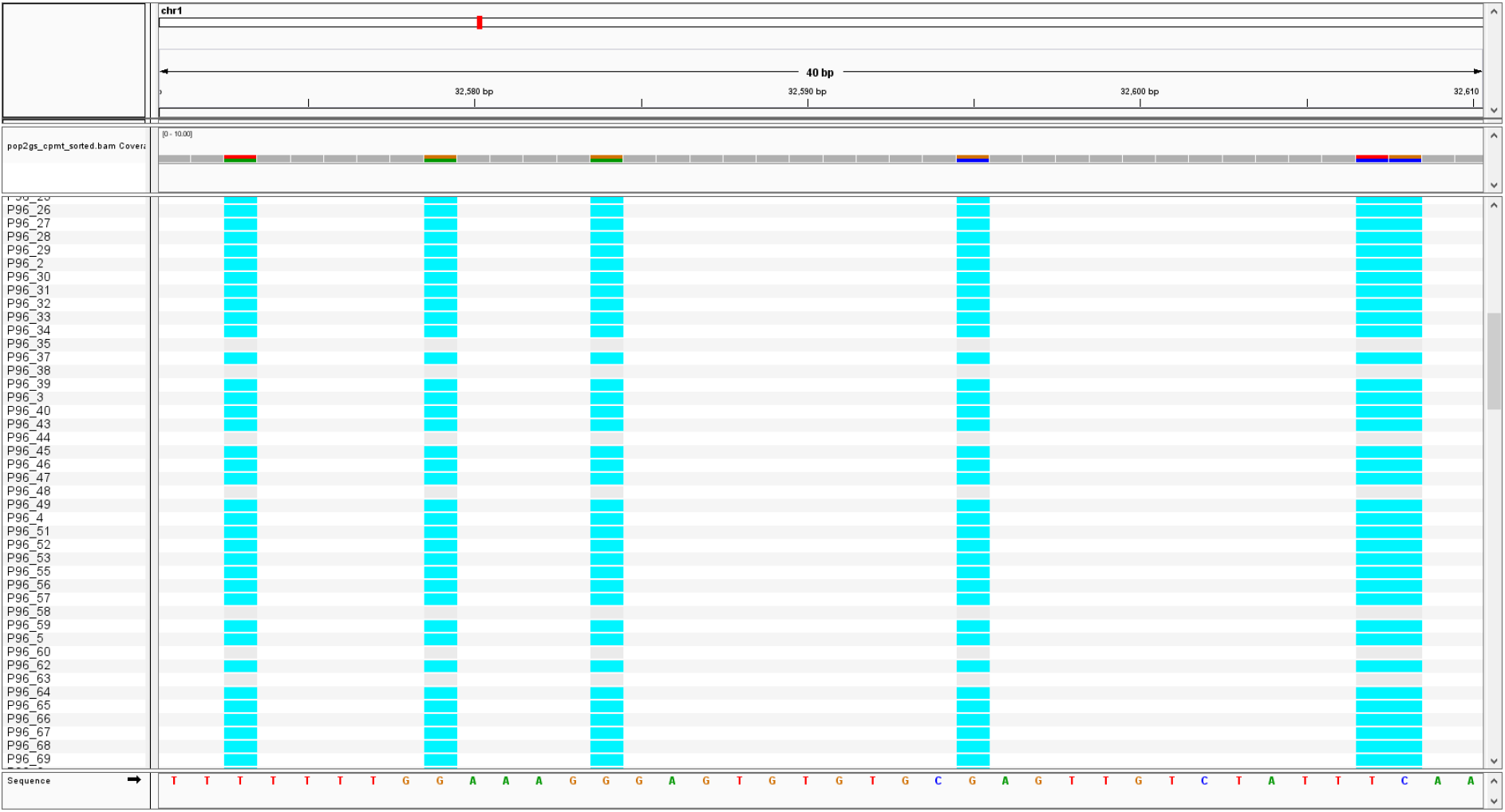
SNP variation among samples of ryegrass population P96, as shown in a region of cpDNA where six SNPs are called; plants P96-35, 38, 44, 48, 58, 60, 63 exhibit different SNP variations compared to the other plants. For example, P96-35 (gray) are the same as the reference at all six SNP sites, while P96-37 (blue) has different site genotypes from the reference.

With an interest to identify SNPs that are differentially distributed between the two populations we performed statistical testing with the R function “prop.test”. There were 99 SNPs identified as significant (Benjamin & Hochberg adjusted p-value < 0.01), out of which 15 were cpSNPs and 84 mtSNPs. A detailed analysis was conducted as follows regarding the correlation and origin of these significant SNPs.

### Correlation and origin of cpSNPs

Comparing to Hamming distance the phi coefficient has an intuitive interpretation as Pearson’s correlation coefficient, in that the coefficient is ranged from −1 (perfect negative association) and 1 (perfect positive association). We performed a hierarchical clustering analysis of the 15 cpSNPs based on phi coefficient (1-ϕ), and the result was shown in Fig. 3. Two negatively correlated groups were largely formed for cpSNPs. In general, adjacent cpSNPs (the SNP location is encoded in the ID), such as cp68818 and cp68834, cp48913 and cp48940 were perfectly correlated, a trend often observed for nuclear SNPs (nuSNPs). However, cp77036 and cp77039 were adjacent SNPs but negatively correlated (ϕ = −1) across samples of the two populations. This revealed a fact that plants with number genotype 0 (see Methods) at cp77036 must have genotype 1 at cp77039 (Table 1).

**Fig. 3.**
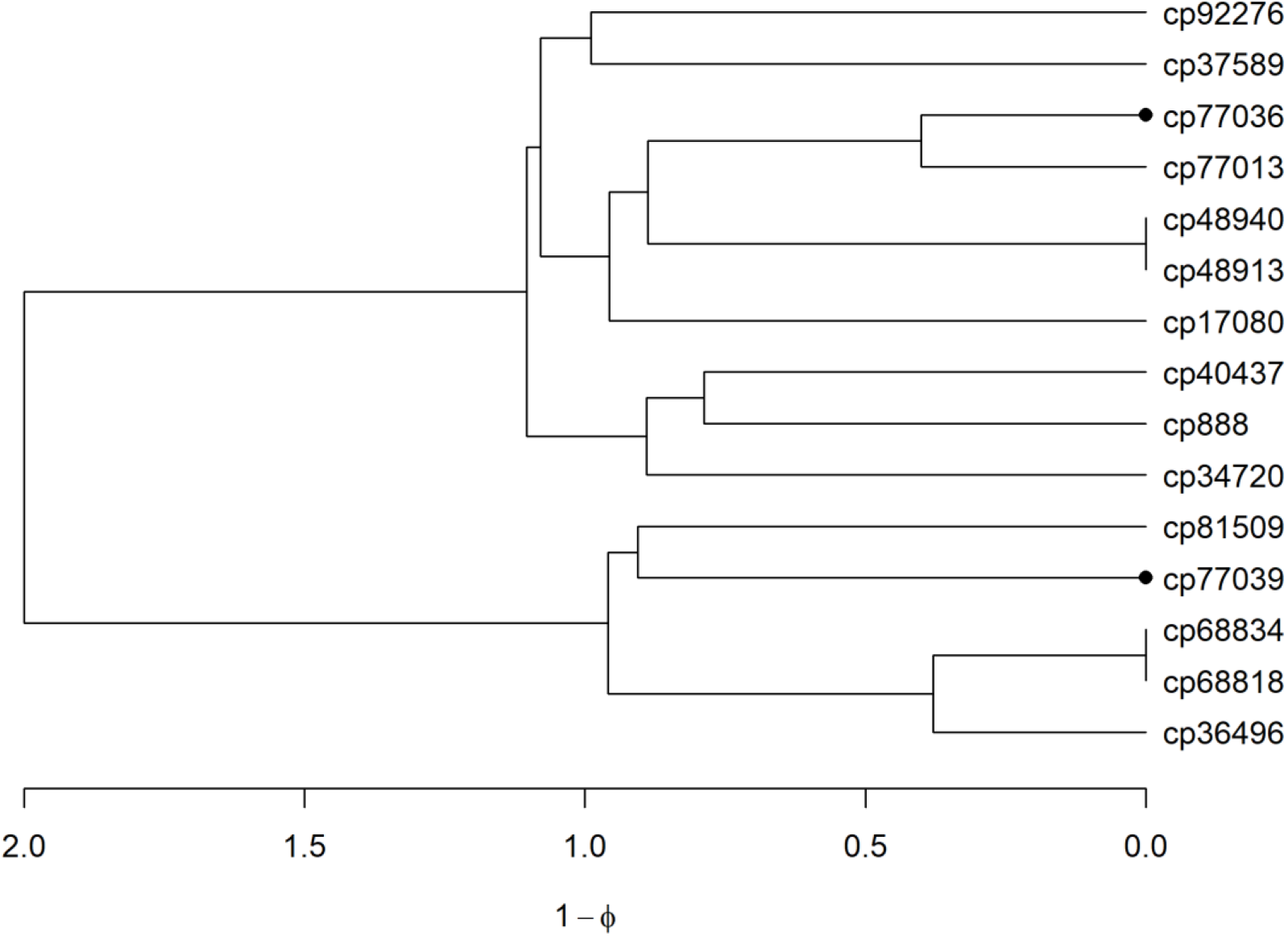
Hierarchical clustering of 15 cpSNPs based on distance metric (1-ϕ). Two SNPs (cp77036 and cp77039) as dotted are adjacent but negatively correlated in contrast with other adjacent SNPs.

**Table 1.**
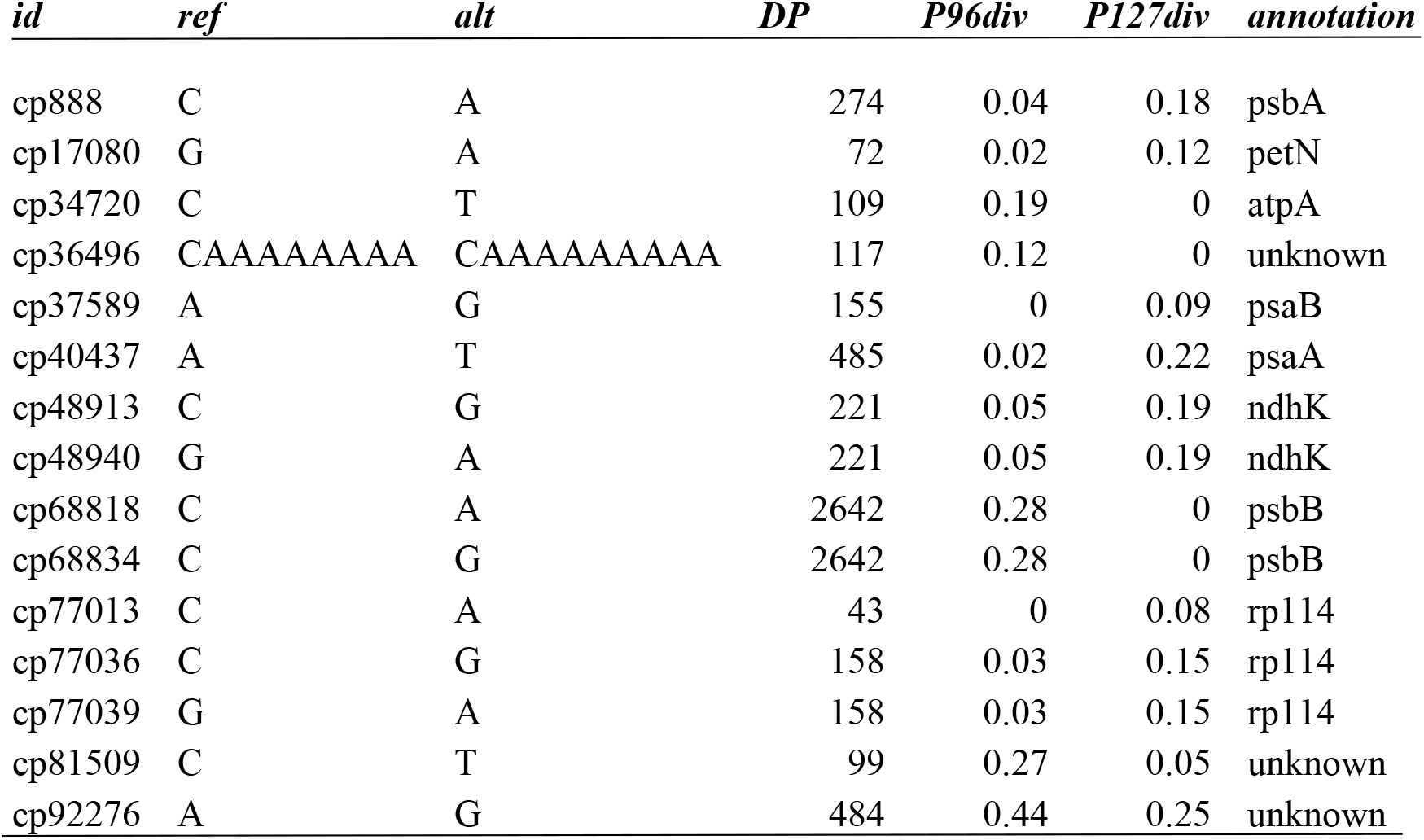
Population diversity and annotation of cpSNPs. A summary of 15 cpSNPs including column information: ***id***, SNP id with prefix cp denoting chloroplast origin, and integer denoting its physical position in the chloroplast genome; ***ref***, reference site base; ***alt***, detected alternative base. ***DP***, read depth; ***P96div***, SNP genotype may be population-dependent, for example, cp34720 has genotype C > T in 19% of plants in the population P96, as shown P96div score = 0.19; ***P127div***, this SNP cp34720 has only genotype C > C in all P127 plants, with P127div score = 0. cp36496 is an indel also presenting as population-specific; ***annotation***, gene symbols, with the full name listed in the Abbreviation.

It was our observation that some SNP genotypic variation was population-dependent. For example, cp68818 had genotype C > C, (a letter genotype denoting ref base > site base, see Methods) for all ‘P127’ plants. However, this SNP showed differentiations in ‘P96’, with 81 plants genotyped as C > C (0) and 31 plants genotyped as C > A (1).

To quantify such SNP variations within a population we defined SNP diversity score as:

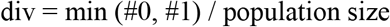

For each population, #0 refers to the number of plants with genotype 0 and #1 the number of plants with genotype 1. For example, for cp68834, its div score in ‘P96’ (P96div) was 0.28 (31/112) while in “P127” the div score = 0. P127div = 0 indicated no differentiation within that population. The maximum SNP div score in a population is 0.5. The div scores for the top 15 cpSNPs were provided in Table 1. Although no difference on average in the diversity score across all 15 cpSNPs (P96div = 0.12, P127div=0.11) there were obvious differences between populations at each individual SNP, which manifested in the population substructures observed in Fig. 1a. Gene psbB (see Abbreviations for the gene symbol description) showed a strong differentiation in ‘ P96’ only, while psbA, ndhk and rpl14 genes were more diverse in ‘P127’. An indel (cp36496), located upstream of gene psaB (36513 – 38717), was differentiated in ‘P96’ only. cp92276 of unannotated origin exhibited a high degree of variation in both populations.

### Correlation and origin of mtSNPs

Likewise, we conducted the investigations on mtSNP diversity within the populations. The clustering of the 84 significant mtSNPs (Supplementary Fig. 2) presented some redundant information in that the adjacent SNPs were highly, positively correlated. For simplicity and without the loss of information, we merged the highly correlated (ϕ > 0.9) and adjacent (< 64 bp) SNPs for discussion. The remained 57 mtSNPs formed two distinct groups (Fig. 4). One group contains 21 mtSNPs which were characterized as P127-diverse (div scores, mean = 0.11, sd = 0.08), i.e. homogeneous in ‘P96’ (mean = 0.05, sd = 0.06). Another group of 36 mtSNPs is a P96-diverse group with P96div (mean = 0.15, sd = 0.096) and P127div (mean = 0.05, sd = 0.09).

**Fig. 4.**
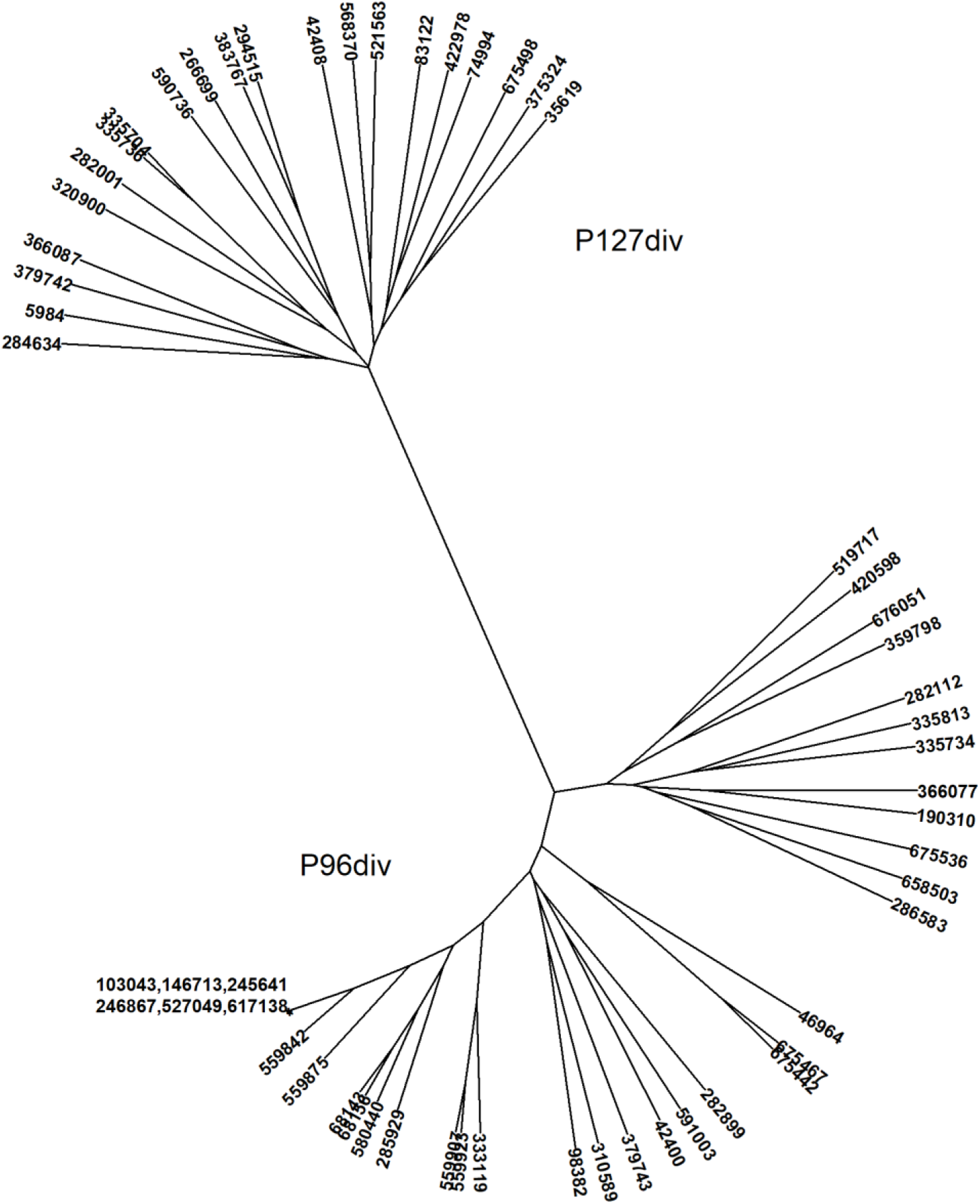
Hierarchical clustering mtSNPs based on 1-ϕ. Two sets of SNPs are shown with the diverse scores distinct between P127 and P96. More differentiations in SNP variation are observed in P96. *: 6 mtSNPs with ϕ = 1 but not adjacent.

Summarized in Table 2 included the diversity score and annotation of the 57 mtSNPs. Annotations for the mtSNPs are scarce due to the fact that about 79% of the ryegrass mitochondrial genome sequences have not been annotated (Islam et al. 2013). In addition to the population-specific mtSNP variations we also made the following observations: (1) For adjacent SNPs, although highly positive correlation was the main trend, the variations in the level and even direction of correlation did occur: from ϕ = −1 for pairs such as mt42400/42408 and mt366077/366087, to low correlation pairs such as mt379742/37974 (ϕ = −0.04), mt335734/335736 (ϕ = −0.13) and mt675467/675498 (ϕ = 0.08); (2) SNPs derived from rrn18 gene were heterogeneous in ‘P96’ but remained homogeneous in ‘P127’, showing population-specific type of variation; in contrast, SNPs from atp4 appeared to be random between the populations; (3) The most variable region in the mitochondrial genome were in the rrn18 gene, within which both rrn5 and ccmFC are resided; (4) 19 mtSNPs (including adjacent and correlated SNPs) were found from position 580,440 to 676,066 bp, indicating another variable but unannotated region. We performed a RepeatMasker analysis (with *Poacea* database)(Tarailo-Graovac and Chen 2009; Bao et al. 2015), and found that this region hosts a number of microsatellite sequences and LTR/Gypsy (Supplementary data 2). (5) Last but not least is the low SNP diversity scores (as also relevant for Table 1). It deserves a further investigation on whether the low div scores e.g. < 0.05 are considered as the noise level.

**Table 2.**
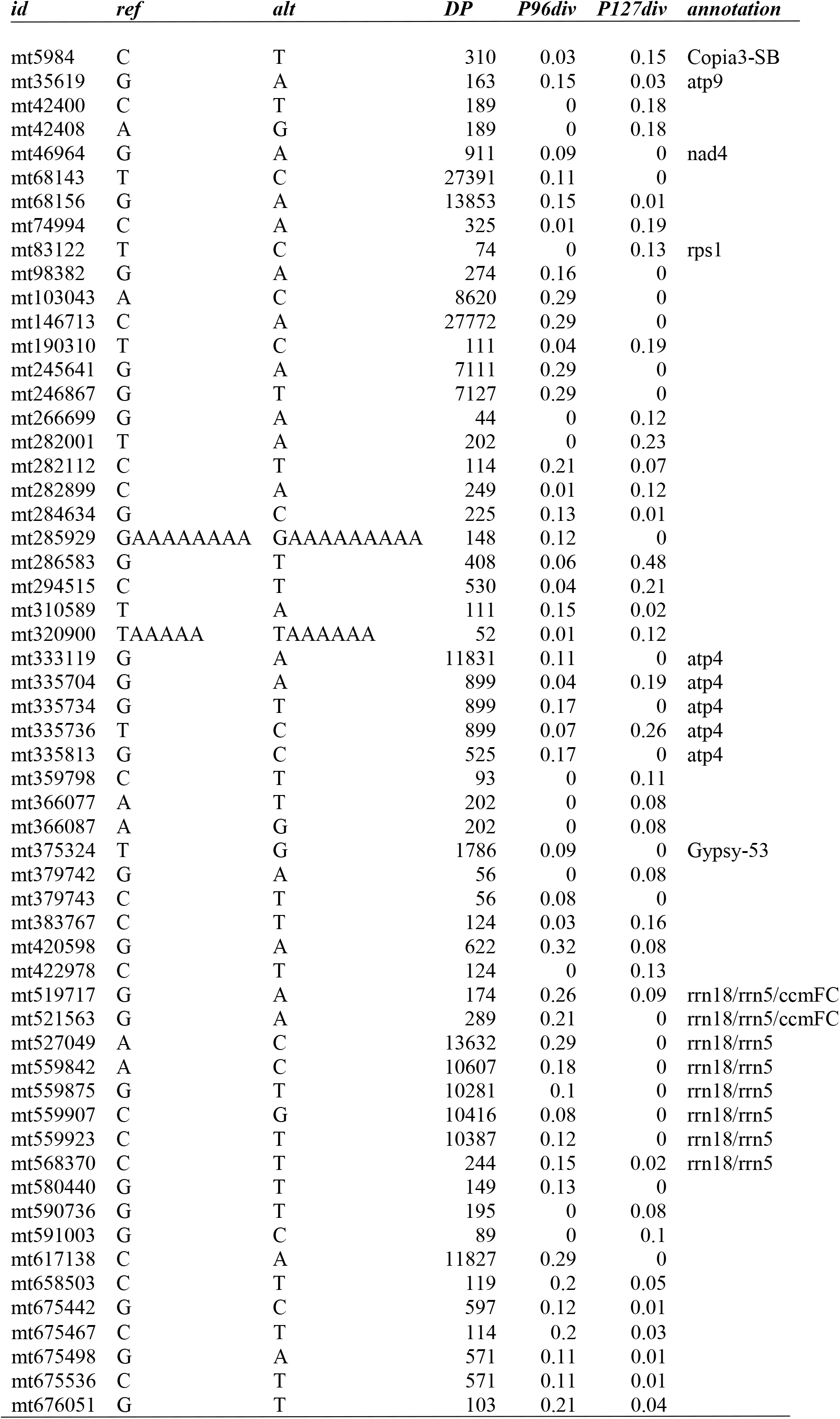
Population diversity and annotation of mtSNPs. A summary of 57 mtSNPs including same column information as described in Table 1. Here, SNP id with prefix “mt” denoting mitochondrial origin. Gene rrn5 is located inside of rrn18 denoted as rrn18/rrn5.

Genetically, both ‘P96’ and ‘P127’ have undergone complex breeding crossing history. As is typical in breeding outcrossing perennial ryegrass, ‘P96’ was initiated as a cross between two breeding populations. This was achieved by crossing 18 pairs of individuals selected randomly from the respective populations. Seeds were harvested both parental populations and advanced through to F2 population, and after cycles of selection to form ‘P96’. Therefore, ‘P96’ is cytoplasmic heterogeneous, with cytoplasm derived from the both breeding populations. While ‘P127’ was started as an inter-cross amongst five cultivar populations, but only seeds were harvested from one of parental cultivars and advanced to form ‘P127’. So, the cytoplasm of ‘P127’ was only derived from a single cultivar. This largely explains the result that “P96” was more diverse than ‘P127’, revealed by mtSNPs (Fig. 1b). The overall diversity scores based on the mtSNPs (P96div = 0.15, P127div = 0.11, Fig. 4) indicated higher degree of diversity indeed occurred within ‘P96’ than that within ‘P127’.

However, the population structures revealed by cpSNPs (Fig. 1a) remained unexplained with our current knowledge. Plastid (chloroplast or its precursor) inheritance is more intriguing. Among flowering plants, biparental inheritance has been reported more common in plastids than in mitochondria (Mogensen 1996), and chloroplast biparental inheritance was perceived as more common in outcrossing species (Reboud and Zeyl 1994).

### Cytoplasmic markers identified from a PstI-MspI GBS library

GBS uses restriction enzymes for genome complexity reduction so there are many modifications in the choice of enzymes. The use of a two-enzyme PstI-MspI GBS protocol (Poland et al. 2012) further reduces genome complexity and increases read depth for more reliably calling of nuSNPs. It was therefore of interest to evaluate how this double-enzyme GBS protocol would affect cytoplasmic SNP marker discovery. To this end, a dataset based on another two ryegrass populations, ‘S96’ and ‘S127’, was recruited. From the two respective populations 45 and 49 plants were sampled. All the samples underwent the PstI-MspI digestion in one GBS library construction.

Out of 1,545,311 tags 1,038 unique ones were mapped to ryegrass cpDNA and 4,631 to mtDNA. In total, 337 SNPs were identified via the same procedures as applied to the ApeKI libraries. Of these 337 SNPs 65.9% SNPs (222) were uniformly distributed between ‘S96’ and ‘S127’, varied only with the corresponding reference sites. After removal of the uniform SNPs 115 SNPs (23 cpSNPs and 92 mtSNPs) were retained for statistical analysis. The similar results were obtained as that from the ApeKI data. MDS analysis based on cpSNPs (Fig. 5a) showed no overall difference between populations, but discernible within-population variation. And the variations due to mtSNPs largely explained the two populations (Fig. 5b).

**Fig. 5.**
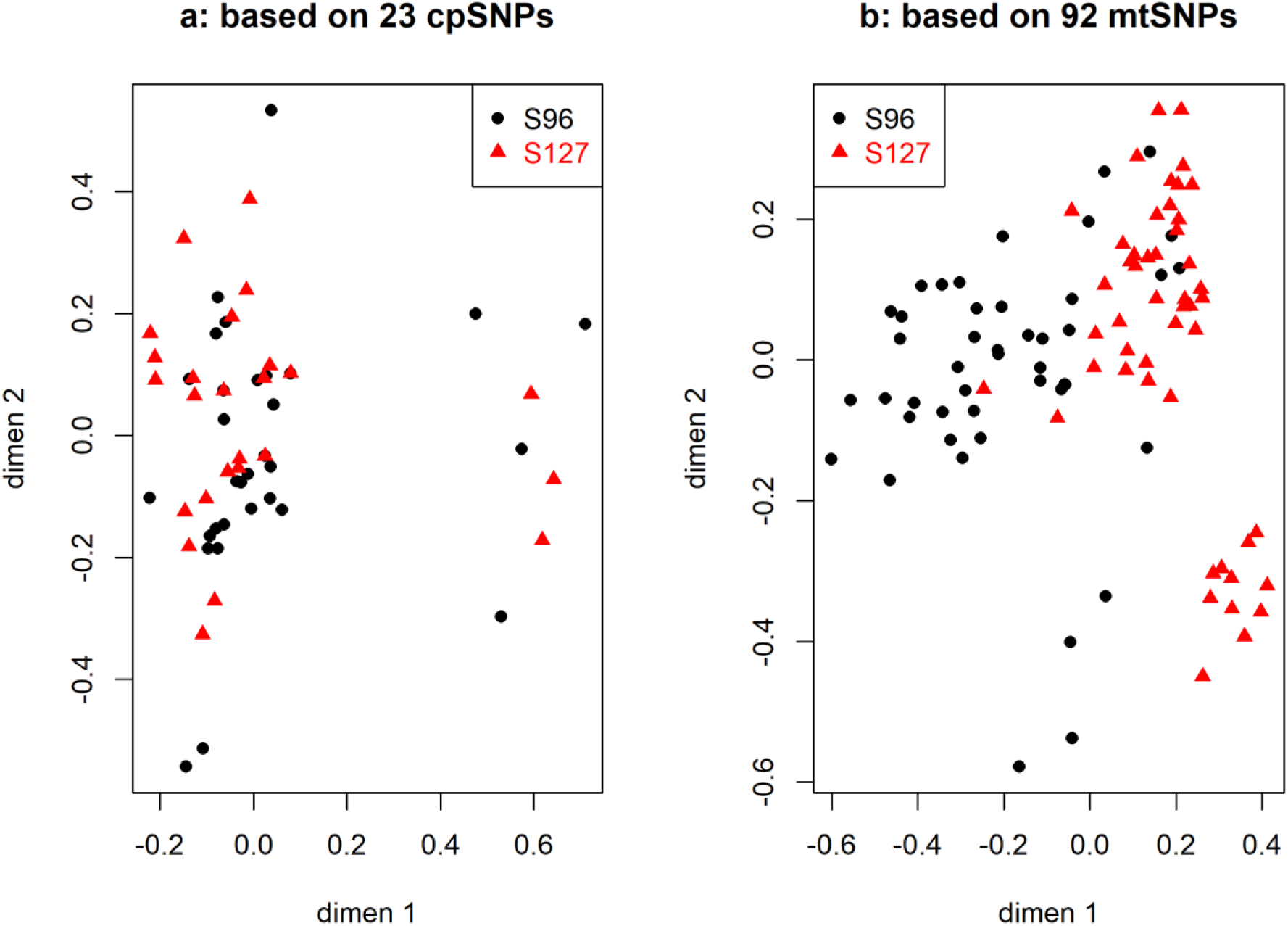
Multidimensional scaling (MDS) clustering of samples based on cpSNPs (a) and mtSNPs (b), from a PstI-MspI library.

Based on the same criteria we used for ApeKI data (BH adjusted p-value < 0.01) only 12 SNPs showed significantly different between ‘S96’ and ‘S127’ (Table 3), among which adjacent SNPs (cp60150/60167, mt636996/636998, mt664895/664898/664924) were all perfectly correlated (ϕ = 1).

**Table 3.**
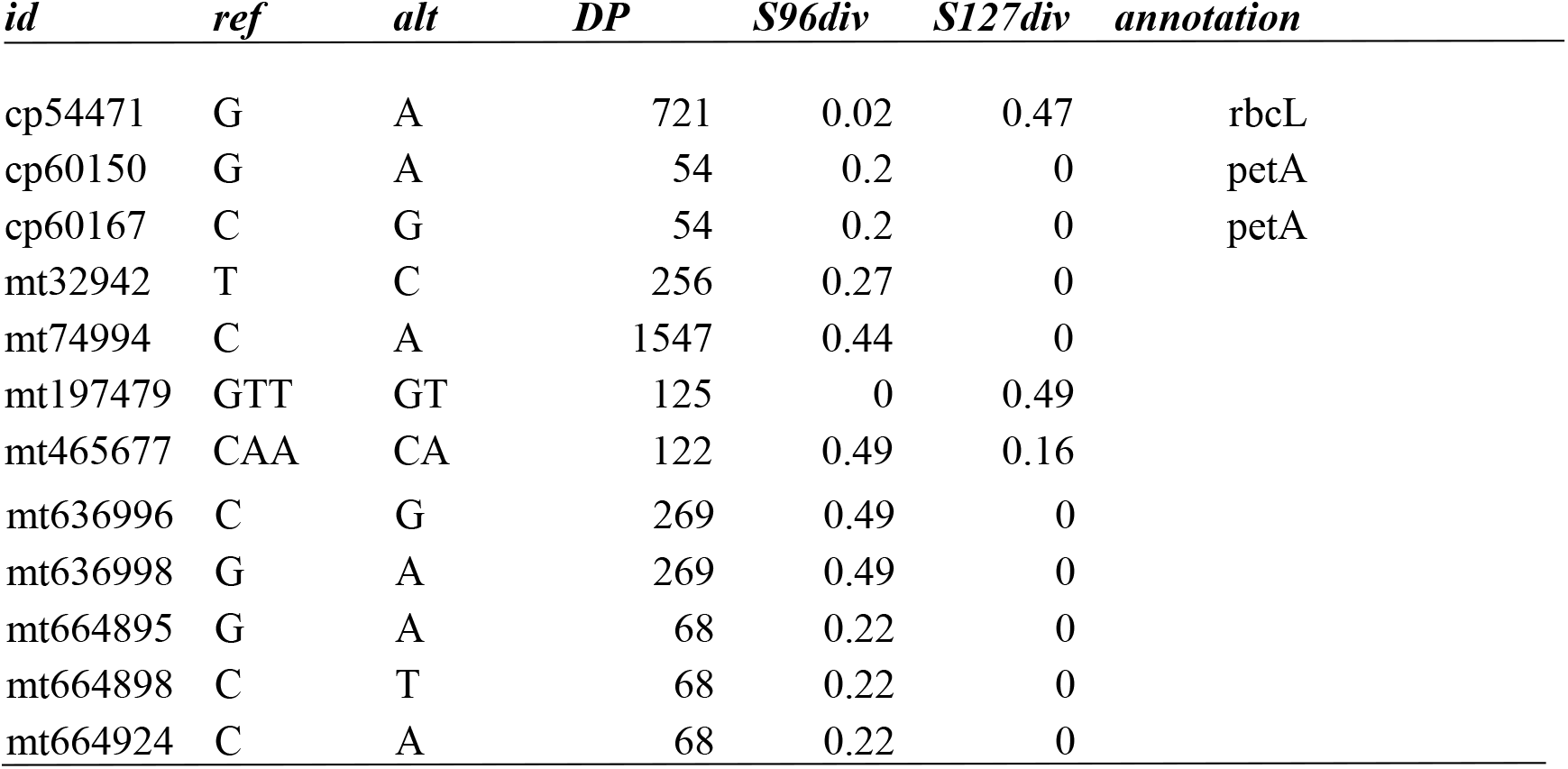
SNPs called from a PstI-MspI GBS library. A summary of SNPs including the same column information as described in Table 1 and 2.

| *id* | *ref* | *alt* | *DP* | *S96div* | *S127div* | *annotation* |
| --- | --- | --- | --- | --- | --- | --- |
| cp54471 | G | A | 721 | 0.02 | 0.47 | rbcL |
| cp60150 | G | A | 54 | 0.2 | 0 | petA |
| cp60167 | C | G | 54 | 0.2 | 0 | petA |
| mt32942 | T | C | 256 | 0.27 | 0 |  |
| mt74994 | C | A | 1547 | 0.44 | 0 |  |
| mt197479 | GTT | GT | 125 | 0 | 0.49 |  |
| mt465677 | CAA | CA | 122 | 0.49 | 0.16 |  |

|  |  |  |  |  |  |
| --- | --- | --- | --- | --- | --- |
| mt636996 | C | G | 269 | 0.49 | 0 |
| mt636998 | G | A | 269 | 0.49 | 0 |
| mt664895 | G | A | 68 | 0.22 | 0 |
| mt664898 | C | T | 68 | 0.22 | 0 |
| mt664924 | C | A | 68 | 0.22 | 0 |

For the three significant cpSNPs (Table 3), cp54471 from gene rbcL remained homogeneous within ‘S96’ but was heterogeneous (S127div = 0.47) in ‘S127’, and cp60150/cp60167 from gene petA (cytochrome f) were diverse only in ‘S96’. All the mtSNPs (Table 3) were derived from the un-annotated regions. mt636996 and mt664895 were derived from a region where several Gypsy transposons are located (Supplementary data 1). We have observed the number of nuSNPs is much less from PstI-MspI library than that from ApeKI library (our unpublished data), and the same was observed here for the cytoplasmic markers.

The 95 samples were generated from one PstI-MspI GBS library and sequenced in two lanes on the same flowcell. This, therefore, confirmed that the sample clustering based on cp or mtSNPs was not confounded by the technical factors of library batch or flowcell.

We have also checked whether the sequencing tags of fungal endophyte origin could possibly affect the observed population structures, as endophytes (*Epichloë* spp.) are known to naturally reside in *L. perenne* cultivar populations. We checked this by mapping all the tags (PstI-MspI library), that have been mapped to ryegrass cytoplasmic genomes, to an endophyte reference genome (*Epichloë festucae*, NCBI Accession PRJNA51625). No single SNP was obtained after following the same SNP calling procedures. Therefore, the endophyte factor hypothesized to influence the observed ryegrass population substructures was ruled out.

## Discussion

We have demonstrated the effective discovery of cytoplasmic genome-wide markers based on genotyping-by-sequencing data in perennial ryegrass. This methodology enables us to characterize large number of samples, to survey gene variant right across cytoplasmic genomes, and to monitor within-population variations of chloroplast or mitochondrial origin.

Our approach depends on the assumption that sequencing tags derived from cpDNA and mtDNA must be available along with that from nuDNA, and the number of tags should be sufficiently large for reliable SNP detection. Provided hundreds and thousands of chloroplasts and mitochondria exist in each plant cell (Cole 2016), that assumption has also been proved true in this study. We have examined and then removed several possible factors, including GBS libraries, sequencing batches and endophyte, which might affect the observed within-population structures. Nonetheless, we recognize, compared to that revealed by nuSNPs (Supplementary Fig. 3), the within-population variation is of a different pattern, and more subtle substructures occurred. Robust modelling of noise level (e.g. to determine the threshold of SNP diversity score) should enhance signals and improve the estimate of accurate biological variations within a population. On the other hand, for a certain ryegrass population, the cytoplasm of its maternal population may still be of a mixed origin (with seeds harvested from both parents in earlier generations), which may contribute the observed complex patterns. It lacks methods to characterize systematically cytoplasmic genetic changes over many generations. Our methods now pave a way to design experiments to monitor cytoplasmic genetic changes in response to different breeding crossing schema, and to quantify the strength of biparental inheritance of chloroplast in ryegrass and other plants.

Maternal inheritance of cytoplasmic genes is widely accepted, but the exact mode of action varies from species to species (Reboud and Zeyl 1994), and largely remains to be elucidated (Birky 2001; Luo et al. 2018). CMS (cytoplasmic male sterility) gene and its use in crop production have been widely-studied in plants (Chase 2007; McDermott et al. 2008), and served as a model for the study of cytoplasmic-nuclear interaction. However, the strength of cytoplasmic-nuclear interaction (Dobler et al. 2014) and co-evolution of cytonuclear integration (Sloan et al. 2018) need to be assessed accurately. Pertaining to plant improvement, sequence variation of cytoplasmic genomes has been found associated with adaptability (Galtier et al. 2009; Bock et al. 2014). Markers derived from mtDNA and cpDNA were found polymorphic among barley cultivars (Hisano et al. 2016), or among individual plants of the same cultivar (Diekmann et al. 2009; Diekmann et al. 2012), but with a few plants genotyped. Cytoplasmic-nuclear interactions are under-explored in plant breeding. Cytoplasmic genome-wide markers as we developed here will facilitate the further understanding of cytoplasmic genome-wide variation by conducting systematic investigations of cytonuclear interactions. This can be achieved without the limitation of the models based on a few genes and a few samples.

In conclusion, we have demonstrated that cytoplasmic markers are readily identified from existing genotyping-by-sequencing (GBS) workflow. The cytoplasmic genome-wide markers contribute a new dimension of genetic information for characterizing breeding populations. Analyses based on markers from chloroplasts and mitochondria reveal different population substructures, providing realizable opportunities to study its mode of action upon selections. Further improvement will be made in assessing sensitivity in marker discovery and accuracy in marker utilization. Our methods invite in-depth exploration of the implications and applications of cytoplasmic inheritance in many other plant species.

## Materials and methods

### GBS data from perennial ryegrass populations

Genotyping-by-sequencing (GBS) have been conducted in our lab for numerous perennial ryegrass populations including elite breeding lines and cultivars. GBS library construction, DNA sequencing and data generation were described in a previous publication (Faville et al. 2018). Raw sequencing read data from four ryegrass populations were selected in this study, with two (designated as ‘P96’ and ‘P127’) having sequence data from ApeKI GBS libraries, and two (‘S96’ and ‘S127’) from a PstI-MspI library. Populations ‘P96’ and ‘P127’ consisted of 112 and 120 individuals, respectively, while ‘S96’ and ‘S127’ contained 45 and 49 individuals, respectively. GBS libraries comprising these individuals were sequenced twice and reads from the two sequencing lanes were combined.

### Tag sequence processing, alignment and SNP calling

Sequencing was conducted on HiSeq2000 as 101 bp single-end reads. Quality reads (having a barcode and a cut site, and no N’s) were processed and de-multiplexed using a Java module “GBSSeqToTagDPPlugin” from the TASSEL-GBS pipeline (Glaubitz et al. 2014). The quality reads (referred to as tags hereafter) of 64 nt were saved in a SQLite database, from which we retrieved tag sequences and tag counts. “TagExportToFastqPlugin” was used to export tag sequences into a fastq file; tag counts in each individual sample were retrieved using “GetTagTaxaDistFromDBPlugin”. There were redundant tags from each sample and we retained the redundant tags for calling SNPs in this report. We found similar population structures were resulted by using the unique set of tags for each sample.

The *Lolium perenne* chloroplast genome (135,282 bp, GenBank: AM777385.2) and mitochondrial genome (678,580 bp, GenBank: JX999996.1) and its associated annotations were downloaded and used as reference genomes. GBS tags were mapped to the reference genomes using “bwa-mem” (Li 2013). Functions from “samtools” (Li et al. 2009) were used for alignment data conversion, indexing and sorting. SNP calling was conducted using “bcftools mpileup” and “bcftools call” (Li 2011) with ploidy = 1 to call site genotype of {A, T, C, G}. Two genotype notations were used here and may be referred to as “letter genotype” and “number genotype”. Letter genotype of one SNP like “G > A” represents “ref base > alt base”, where A is called at the position and different from the reference site (G). And “G > G” means that the site base G is the same as the reference base G. For statistical analysis number genotype was used with, for example, “G > G” is encoded as 0, and “G > A” as 1.

### Statistical analysis

Because of the binary nature of haploid cytoplasmic SNP data Phi correlation coefficient and Hamming distance were employed to calculate the genetic relationships between individual plants, and to measure correlations between SNPs. Multivariate statistics was performed using R functions including “prcomp” for principal component analysis, and “cmdscale” for multi-dimensional scaling analysis. Alignment data interrogation and SNP data analysis were performed based on a number of Bioconductor packages (Lawrence et al. 2013), BioJulia packages (https://github.com/BioJulia) and visualized using IGV (Robinson et al. 2011).

### Code availability

Scripts developed for SNP calling and data manipulation can be accessed from https://github.com/AgResearch/cpmt, under the license of GNU GPLv3.

## Abbreviations

rpl14: ribosomal protein L14
psbA: photosystem II protein D1
psbB: photosystem II CP47 chlorophyll apoprotein
petN: cytochrome b6/f complex subunit N
atpA: ATP synthase CF1 alpha subunit
psaA: photosystem I P700 apoprotein A1
psaB: photosystem I P700 apoprotein A2
ndhK: NADH-plastoquinone oxidoreductase subunit K
atp9: ATP synthase subunits 9
atp4: ATP synthase subunits 4
nad4: NADH dehydrogenase subunit 4L
rps1: ribosomal protein S14
rrn18: 18S ribosomal RNA
rrn5: 5S ribosomal RNA
ccmFC: cytochrome c biogenesis
rbcL: ribulose-1,5-bisphosphate carboxylase/oxygenase large subunit
petA: cytochrome f

## Author contributions

MC conceived the scope of this study, developed the computing workflow and conducted data analysis. MC, MF, JJ and MC (Carena) conducted data interpretation and contributed to writing of the paper.

## Acknowledgements

We thank Tom Lyons and Zulfi Jahufer for providing information on the breeding background of the ryegrass populations used in the study; Dan Sun and Alan McCulloch for maintaining HPC (high-performance computing) environment; and Brent Barrett for useful comments. We thank Grasslands Innovation Ltd, New Zealand for accessing to plant materials and genotyping-by-sequencing raw read data.

