## Supplementary Figs and Data for "Genotyping-by-sequencing based cytoplasmic markers underpin population structural variation of perennial ryegrass"

**Supplementary Fig. 1**: Sample clustering of population "P96" (n=112) indicating 3 sub-clusters based on cpDNA. As shown in Fig. 2 plants 35, 38, 44, 48, 58, 60, 63 belongs to the same cluster (3) as the same pattern of variation (6 SNPs as plotted) is presented.

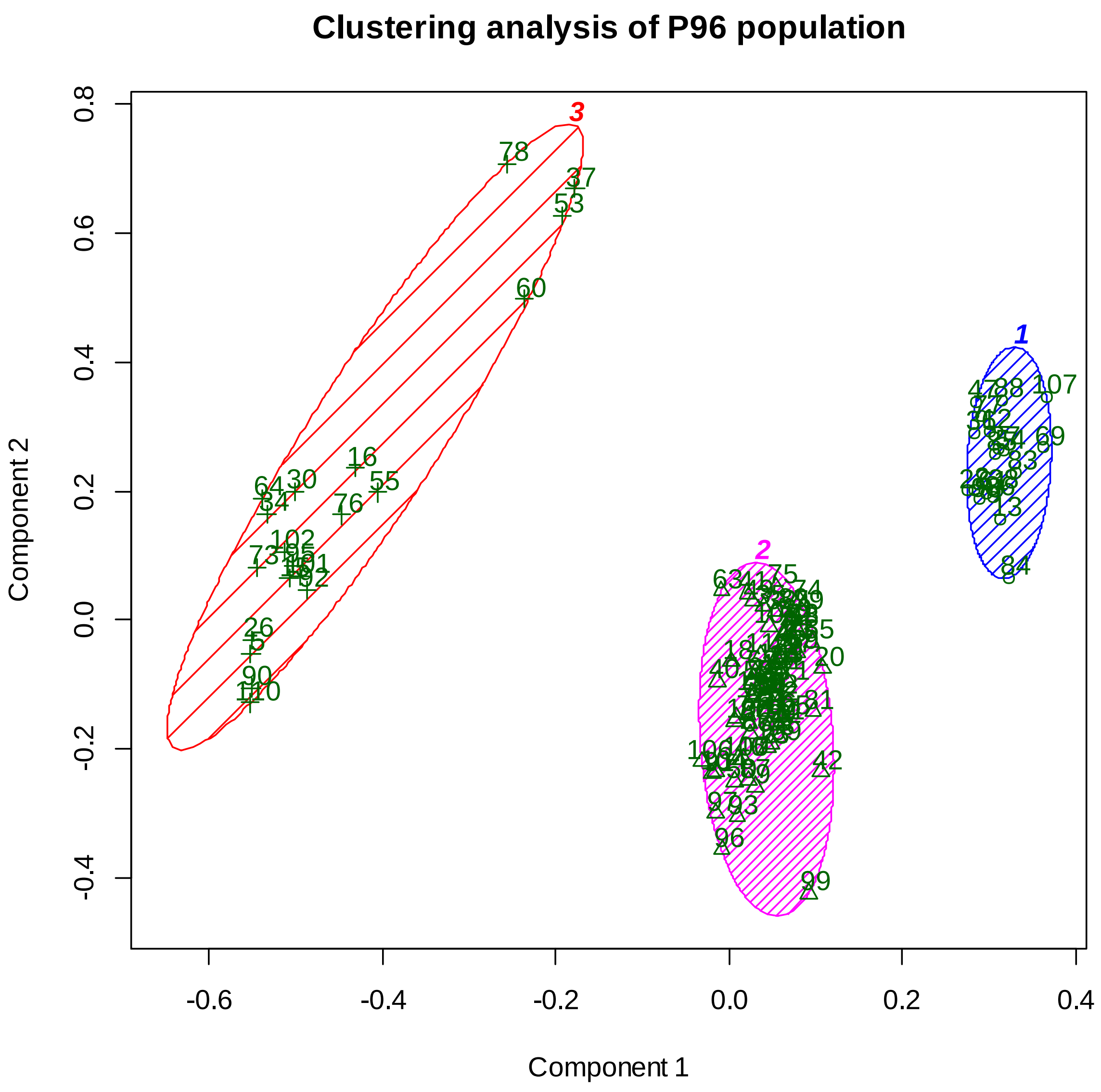

```
> names(tmp)[tmp==3]
 [1] "P96_104" "P96_116" "P96_117" "P96_129" "P96_13"
 [6] "P96_18"  "P96_20"  "P96_35"  "P96_38"  "P96_44"
[11] "P96_48"  "P96_58"  "P96_60"  "P96_63"  "P96_76"
[16] "P96_78"  "P96_81"  "P96_88"  "P96_89"  "P96_98"
```

**Supplementary Fig. 2**: Clustering of 84 mtSNPs (ApeKI library) including the adjacent and highly correlated.

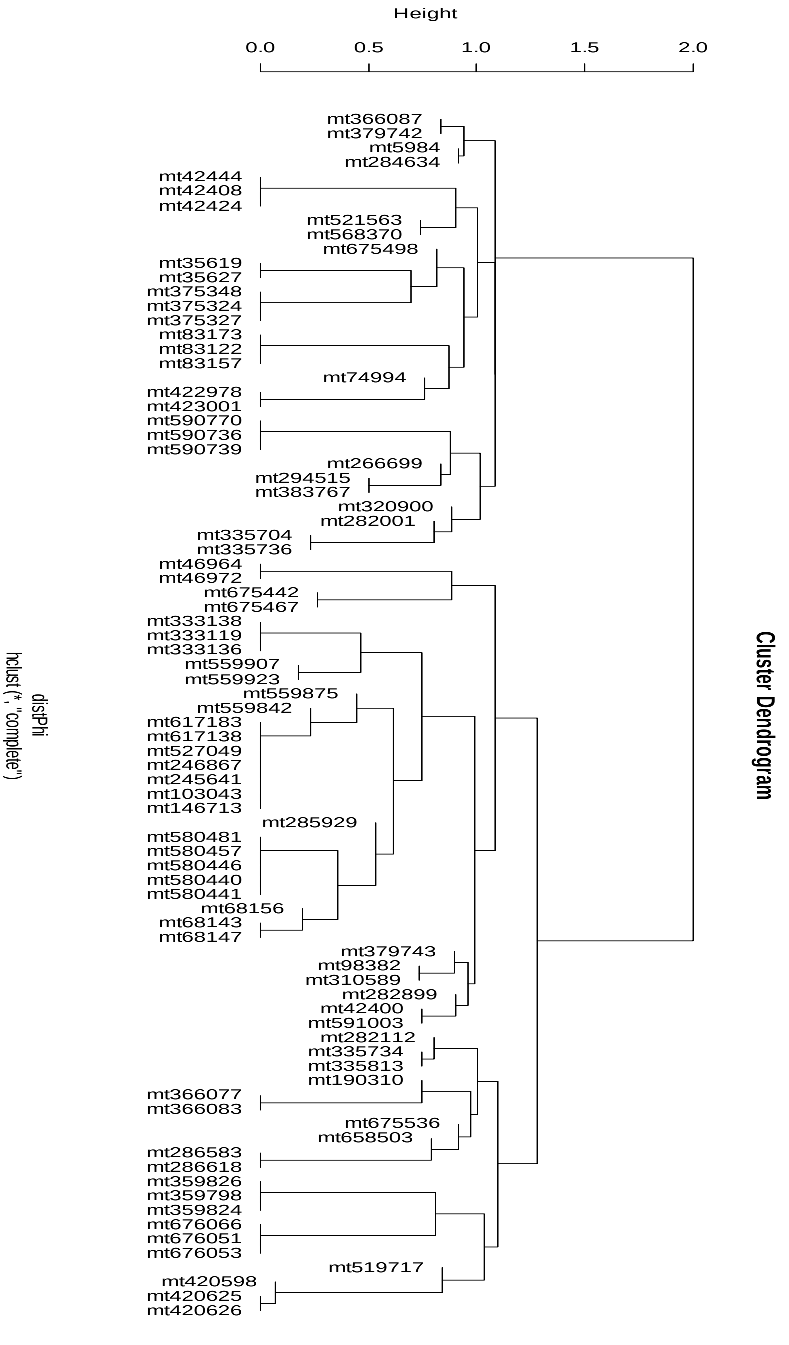

**Supplementary Fig. 3.** A principal component analysis of the two populations ("P96" and "P127") based on 1,223,081 nuSNPs (with filtering of minor allele frequency MAF > 0.05, missingRate < 50% and biallelic only). Tag mapping was conducted using bwa-mem (that used for cytoplasmic marker discovery) to a ryegrass reference genome. The reference genome was described in Faville et al. 2018, where bowtie2 was used for the tag mapping.

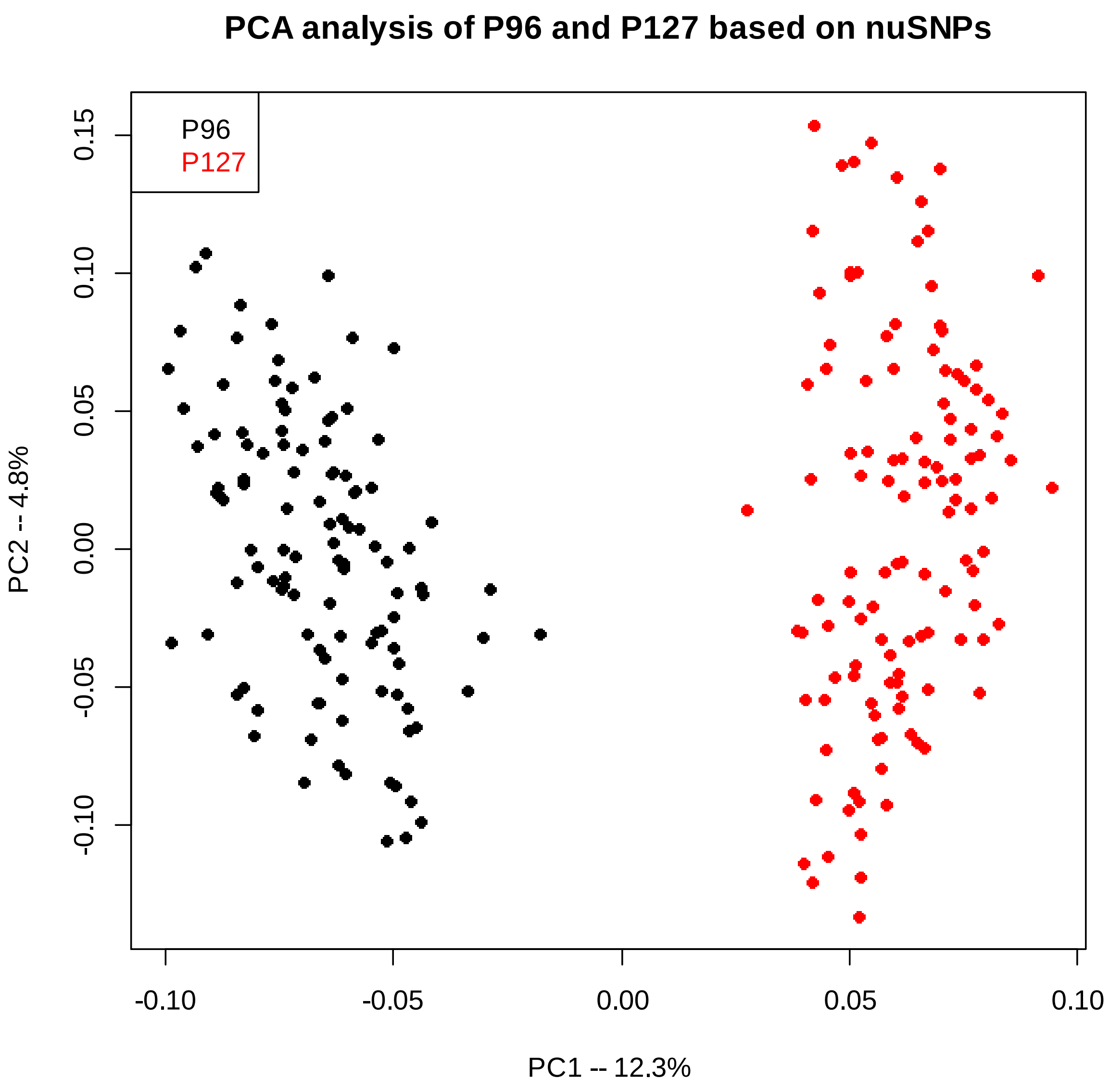


---

**Supplementary data 1**: RepeatMasker annotation of Lp cpDNA, see https://github.com/AgResearch/cpmt/blob/master/data/LpChloroplast_RM.gff3

**Supplementary data 2**: RepeatMasker annotation of Lp mtDNA, see https://github.com/AgResearch/cpmt/blob/master/data/LpMitochondria_RM.gff3
